# Comparison of biochemical properties of RNases HII from *E. coli*, *Geobacillus stearothermophilus* and *Thermus thermophilus*

**DOI:** 10.64898/2025.12.08.693097

**Authors:** Andrei A. Voronin, Igor P. Oscorbin, Lidiya M. Novikova, Mikhail E. Voskoboev, Maxim L. Filipenko

## Abstract

RNases H (EC 3.1.26.4) are a family of enzymes participating in removal of ribonucleotides from double-stranded DNA by hydrolyzing their phosphodiester bonds. Despite a long history of research, several important aspects of RNase HII functioning are poorly known including optimal pH, salts and thermal stability. This lack of empirical data hampers the selection of an optimal RNase HII for a specific practical application. In the present study, we compared biochemical properties of previously cloned RNases HII from *E. coli*, *Geobacillus stearothermophilus* and *Thermus thermophilus*: optimal temperature, pH, salts, divalent cofactors, thermal stability and specific activity. As expected, the most thermostable enzyme was Tth RNase HII, and Mg^2+^ was the most preferential cofactor for all RNases. Gst RNase HII was partially inhibited by K^+^ ions, while other enzymes did not demonstrate preferences for any salt. The enzymes from *E. coli* and *G. stearothermophilus* were typical RNases HII, while Tth RNase HII was a JRNase (junction ribonuclease). All three RNases did not cleave a DNA-RNA_3_-DNA_2_-RNA_1_-DNA/DNA substrate. The presented results will facilitate usage of RNases HII in practical applications and provide a basis for further comparative studies of RNases HII from various organisms.

## Introduction

RNases H are a group of enzymes that hydrolyze phosphodiester bonds of ribonucleotides incorporated into double-stranded DNA. In vivo, they play a key role in removing RNA primers, facilitate the removal of single ribonucleotide monophosphate (rNMP) insertions from dsDNA and are a critical component in resolving R-loops, which leave parts of DNA in a vulnerable single stranded state [1].

RNases H are classified into three types according to their structure and substrate preferences – RNases HI, HII and HIII. RNases HII and HIII are much more closely related structurally, though HI and HIII share greater similarity regarding their substrate specificity [2]. In prokaryotes, RNases HII are usually accompanied by either RNase HI or HIII, but in some archaea, only RNase HII is present. Given this distribution and the rare occurrence of both RNase HI and HIII in a single microorganism, it has been suggested that these enzymes may be functionally redundant due to their similar biochemical properties [3].

Regarding the optimal conditions for RNases H, all three types are dependent on a divalent cation cofactor, such as Mg^2+^, and no detectable activity is observed without them. Although other metal ions have also been tested, the highest specific activity in all studied prokaryotic RNases HII have been achieved with either Mg^2+^ or Mn^2+^. All types of RNases H can generally hydrolyze phosphodiester bonds in substrates such as: RNA/DNA heteroduplexes or R-loop, RNA-DNA/DNA (Okazaki fragment-like substrates) and DNA-RNA_n_-DNA/DNA (if n ≥ 4). Unlike other cognate enzymes, RNases HII can also hydrolyze an additional substrate, DNA-RNA_1_-DNA/DNA. In all substrate types, RNases H hydrolyze phosphodiester bonds to create 3’-OH and 5’-phosphates, although specific hydrolysis sites vary. RNases HII cleave specific sites in various substrates: they cleave the phosphate of the deoxyribonucleotide adjacent to the ribonucleotide in RNA_n+1_-DNA/DNA, producing RNA_n_-3’ and 5’-RNA_1_-DNA; They cleave the sole ribonucleotide phosphodiester bond in DNA_n_-RNA_1_-DNA/DNA, yielding DNA_n_-3’ and 5’-RNA_1_-DNA; and they cleave RNA/DNA in multiple sites (2 to 7 cuts in substrate lengths of 12 to 32 nucleotides), consistently leaving a minimum of 2-3 nucleotides at both ends of the ribonucleotide-containing strand. Cleavage specificity appears to depend on substrate sequence and reaction conditions; however, this has not been thoroughly investigated [1,4–14].

The ability of RNases HII to degrade DNA-RNA_1_-DNA/DNA substrates was once considered unique and is termed junction ribonuclease (JRNase) activity [5,14]. However, it has since been demonstrated that some RNases HI and HIII can also cleave substrates containing a single rNMP under specific conditions. For example, Cp RNase HIII showed a second hydrolysis site, generating DNA_n_-RNA_1_-3’ and 5’-DNA rather than DNA-3’ and 5’RNA_1_-DNA [1,15]. Additionally, *Sulfolobus tokodaii* RNase HII was able to cleave both phosphates of the junction rNMP and junction dNMP in an RNA_9_-DNA_9_/DNA substrate [16].

Each type of RNase’s H activity has practical application. RNA_n_-DNA/DNA oligonucleotides can be probed by RNases H to study RNA primers removal in Okazaki fragments [7]. In RNA-seq, RNase H is used for rRNA depletion, where cDNA is synthesized using only rRNA as a template resulting in RNA/DNA duplexes following rRNA hydrolysis by RNase H. Thus, all non-ribosomal RNA molecules remain intact in the treated sample [17]. Similarly, RNases H are applied for a direct measurement of mRNA poly(A) tails [18]. DNA-RNA_n_-DNA/DNA substrates have found several applications: primers containing a single rNMP and a terminal 3’ blocker effectively prevent non-specific amplification in PCR [19], and fluorescently labelled oligonucleotides probes with rNMP serve for monitoring isothermal amplification (most commonly, LAMP) in a real-time mode [20].

Despite extensive research, several important aspects of RNases HII function remain poorly understood [21]. Notably, there is a fundamental gap regarding optimal conditions for RNase HII biochemistry. Most studies report only optimal cofactors and temperature, often omitting data on optimal pH, salts, thermal stability and tolerance to inhibitors. Moreover, inconsistent experimental designs and conditions hinders comprehensive comparisons of RNases HII from different sources. Only a few reports directly compared various RNase HII using a unified methodology [5]. This lack of detailed biochemical data complicates the selection of suitable RNases HII for practical applications.

In the present work, we compared biochemical properties of previously cloned RNases HII from *E. coli*, *Geobacillus stearothermophilus* and *Thermus thermophilus*. Our goal was to evaluate optimal reaction conditions—including temperature, pH, salts, divalent cofactors—as well as to assess thermal stability. This work aims to facilitate the use of RNases HII in practical applications and provide a basis for further studies of RNases HII from diverse organisms.

## 1. Results

### 1.1. Cloning and purification of RNases HII

Here, we planned to compare the biochemical properties of three previously cloned and studied RNases HII from *E. coli, G. stearothermophilus, and T. thermophilus*, a mesophilic, thermophilic and hyperthermophilic bacteria, respectively. The amino acid sequence homology of Eco RNase HII is 43% with Gst RNase HII and 60% and Tth RNase HII; Gst RNase HII shares 41% similarity with Tth RNase HII (Figure 1). RNases HII are relatively small enzymes with a molecular mass of 20–25 kDa. Although previous studies have partially characterized the biochemical activities of these RNase HII, including optimal temperature, thermostability, preferred cofactors and optimal pH, none have comprehensively evaluated all these parameters within a single study, and no investigation of the effects of salt type and concentration on enzyme activity has been conducted. Tth RNase HII is used for rRNA depletion, and Gst RNase HII also has potential practical application, e.g., isothermal amplification. *E. coli* RNase HII has been studied extensively [22,23] and this enzyme was chosen as a control for the present work.

**Figure 1.**
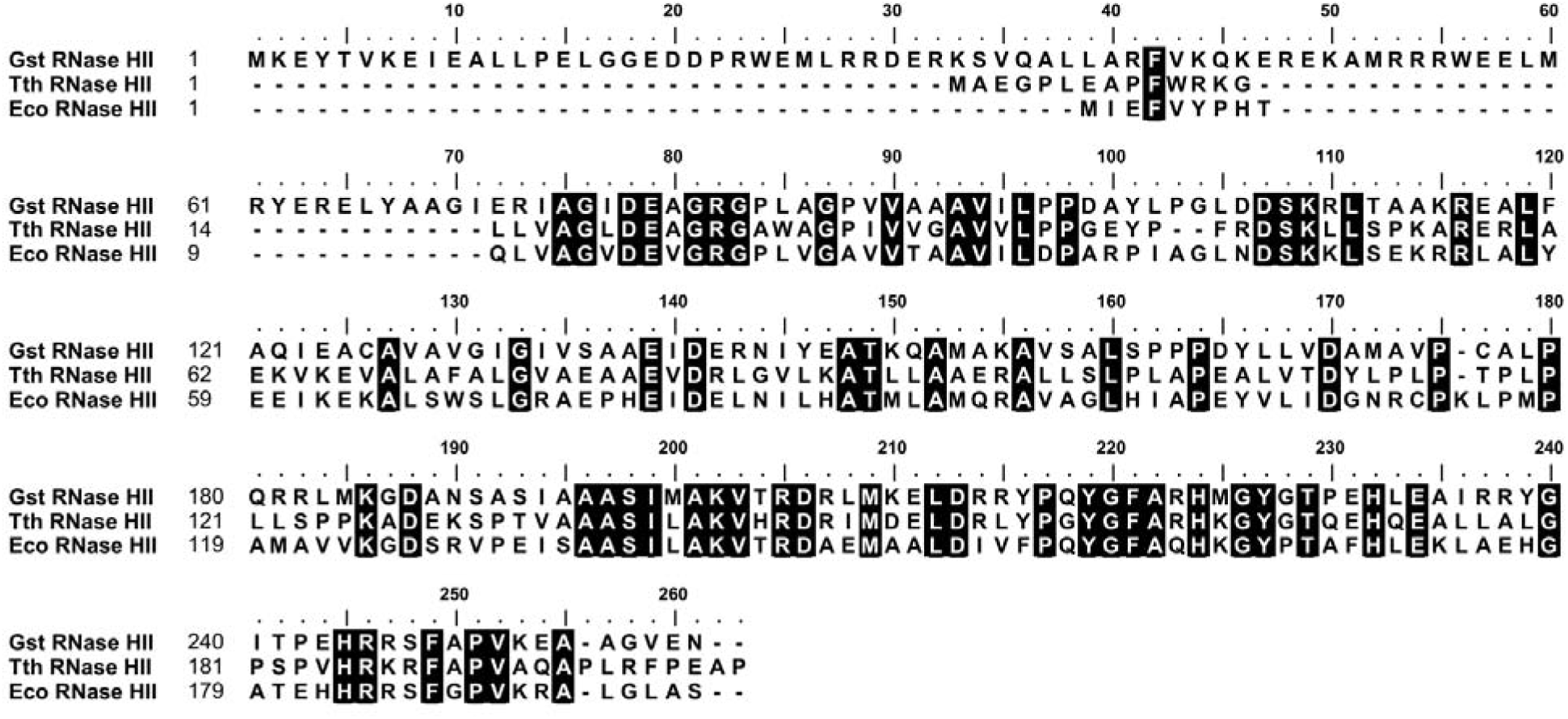
Alignment of RNAse HII amino acid sequences. The GenBank accession numbers are WJQ01363.1 for RNase HII *G. stearothermophilus*, BFH84700.1 for RNase HII *T. thermophilus*, and WP_433794514.1 for RNase HII *E*. *coli*.

As shown in Figure 1, RNase HII of *G. stearothermophilus* possesses an additional N-terminal sequence consisting of approximately 60 amino acid residues. Some RNases HII contain additional sequences at either N- or C-terminus, while others lack any extensions [3]. Although, these additional sequences are apparently redundant for the specific activity of RNases HII, they can enhance enzymatic activity and thermal stability [5].

The studied RNases HII were cloned into pET23d vector (Gst) or pET28a vector (Tth and Eco), expressed in *E. coli*, and purified by metal-chelate and ion exchange chromatography (Figure 2). Electrophoretic purity of RNases was >95%, ∼40% and >90% for Gst, Tth, and Eco RNases HII, respectively. Low electrophoretic purity of Tth RNase HII could be the result of several factors: 1) low expression in *E. coli* despite the usage of Rosetta 2 (DE3) strain with the pRARE2 plasmid, coding tRNAs rare in *E. coli*. 2) low solubility of studied RNases HII that were stable only in the presence of urea or high concentration of NaCl. 3) Tth RNase HII did not bind with cation and anion ion-exchange resins despite changing pH in the range of 6–9. The listed issues will be further thoroughly reviewed in the Discussion section and possible explanations will also be provided. Also, Tth RNase HII could be partially hydrolyzed by *E. coli* proteases due to a possible presence of proteases recognition sites. While these two contaminant proteins could affect the performance of Tth RNase HII, it should be noted that no non-specific nucleases activities were observed for all three RNAses HII. Linear genomic DNA of T7 phage, supercoiled DNA (pQE30 plasmid) and total RNA from human white blood cells from healthy donors remained intact after 24 hours’ incubation at 4°C and 37°C with studied RNAses HII. Concentration of Tth RNAse HII was calculated taking into account its electrophoretic purity to avoid a bias caused by contaminant proteins. Preliminary experiments regarding biochemical properties of Tth RNase HII produced results similar to previously published for this enzyme. Based on these observations, we characterized Tth RNase HII in a manner similar to other RNases.

**Figure 2.**
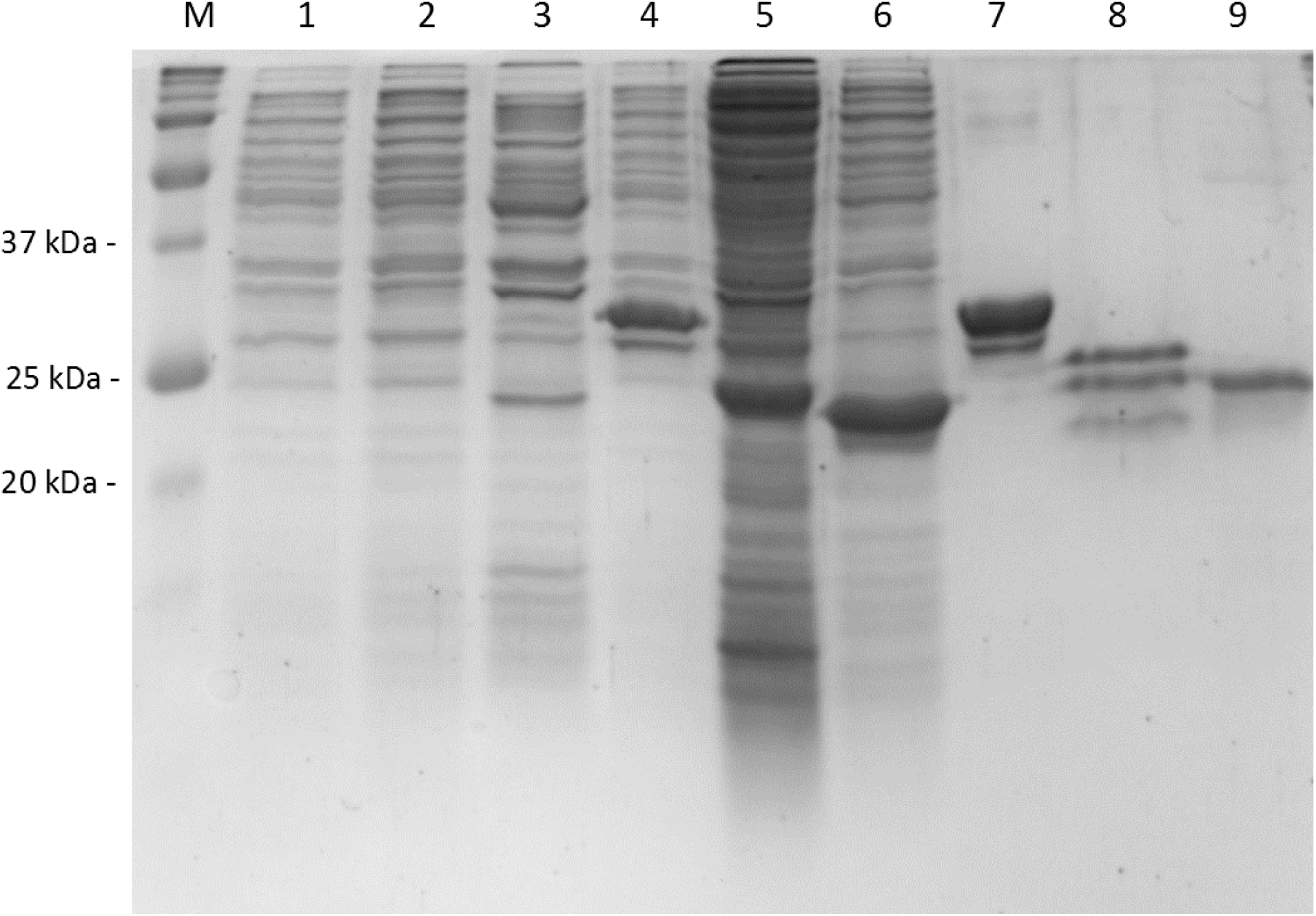
Purification of RNases HII *G. stearothermophilus*, *T. thermophilus*, and *E*. *coli*. Lane 1 — Precision Plus Protein Standard (Bio-Rad, USA). Lanes 2, 3, and 4 — crude lysates before induction; 5, 6, and 7 — crude lysates after induction; 8, 9, and 10 — purified RNases HII. Gst RNase HII— 1, 4, 7; Tth RNase HII— 2, 5, 8; Eco RNase HII— 3, 6, 9.

### 1.2. Optimal temperature and thermostability of RNases HII

For many practical applications, enzymes with high optimal temperatures and thermostability are advantageous. Also, highly thermostable enzymes generally retain significant activity at lower temperatures, making them more versatile [24]. Thus, the optimal temperature for studied RNases HII were measured at the first stage of optimization. RNase HII activity was assessed in a reaction buffer recommended for Eco RNase HII by NEB and a fluorescently labelled duplex DNA substrate (H2-TM3/H2-TM3RT) containing a single rNMP. In the reaction buffer, the substrate has a calculated melting temperature of approximately 75 °C, allowing measurement of RNase HII activity up to 75°C. Preliminary, we analyzed specific activity RNases HII to determine enzymes amounts sufficient to hydrolyze approximately 50 % of the substrate. This titration allowed to avoid reaction saturation. The reaction temperature was varied from 20 to 75°C. The results are presented in Figure 3.

**Figure 3.**
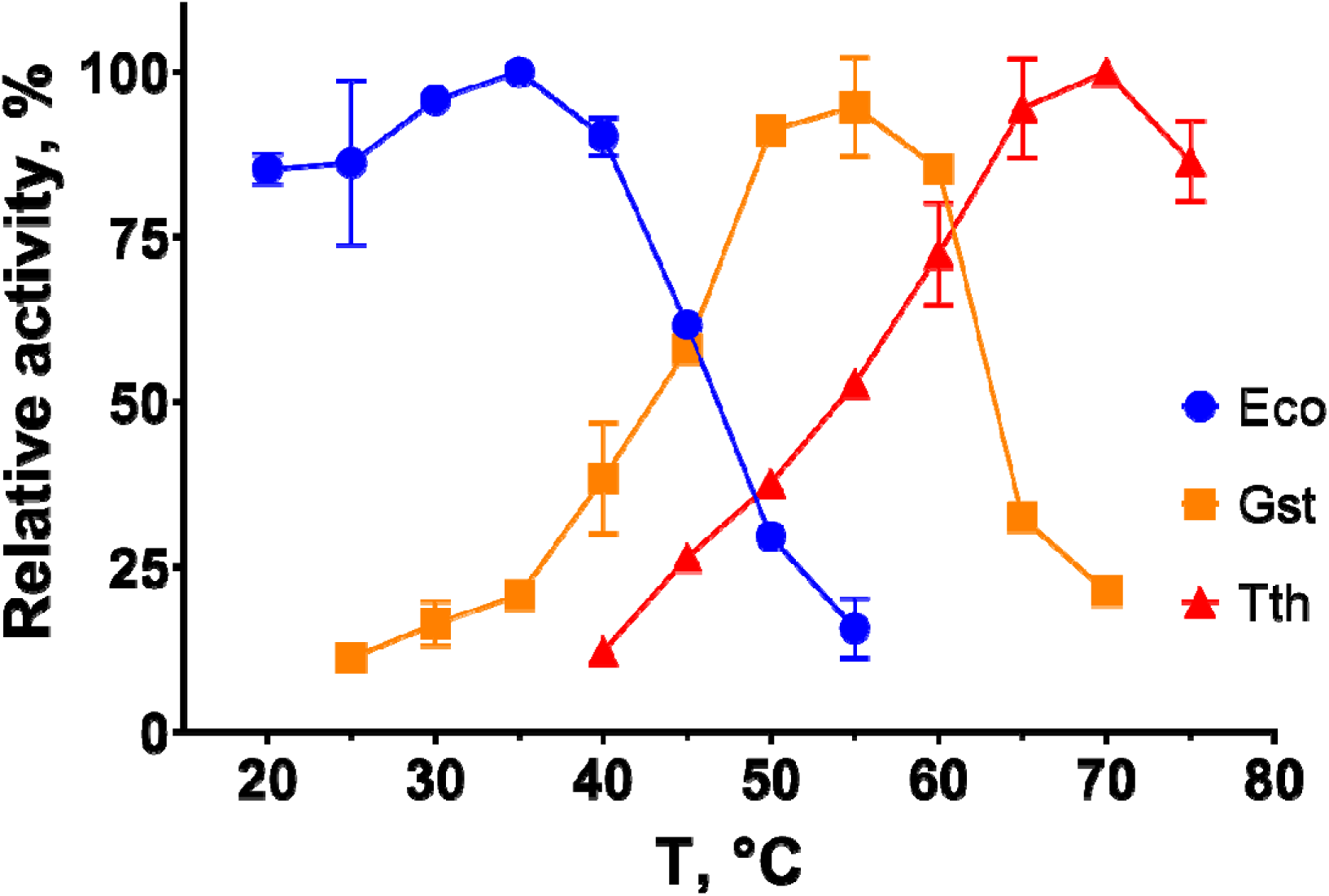
Optimal temperature for RNAses HII *G. stearothermophilus*, *T. thermophilus* and *E. coli*. A duplex DNA oligonucleotide with a single rNMP on a fluorescently-labelled strand served as a substrate. All reactions were conducted in the optimal buffer for Eco RNase HII (NEB) with 0.01 U_1_ of an enzyme (determined as stated in paragraph 2.4). Y-axis represents the relative activity of each RNase HII, X-axis indicates the reaction temperature. Curve colors correspond to each RNase HII: blue for Eco, orange for Gst, red for Tth RNAse HII.

*T. thermophilus* and *G. stearothermophilus* are thermophilic bacteria with an optimal growth temperature higher and close to 60 °C, respectively. Therefore, a high optimal reaction temperature was expected for RNases HII from these organisms. As shown in Figure 3, the optimal temperature for Gst RNase HII was 55 °C, for Tth RNase HII was active 70 °C, and Eco RNase HII was the most active at 35 °C under the assay conditions. All subsequent experiments were conducted at each enzyme’s optimal temperature.

Thermal stability was evaluated by pre-incubating 0.01 U_1_ of RNase HII in a reaction buffer without the substrate, varying the incubation time and temperature, followed by the activity assay described above. Pre-incubation was conducted over a temperature range of 30–80 °С for 5–90 min, and the results are shown in Figure 4. Gst RNase HII retained over 95% of its activity after 90 minutes at 50 °C but rapidly lost activity at 60 °C. Tth RNase HII lost about 30% of its activity after incubation at 70 °C. Eco RNase HII was significantly inactivated by incubation at 40 °C for more than 30 minutes but surprisingly retained about 20% of its activity after 15 minutes at 80 °C.

**Figure 4.**
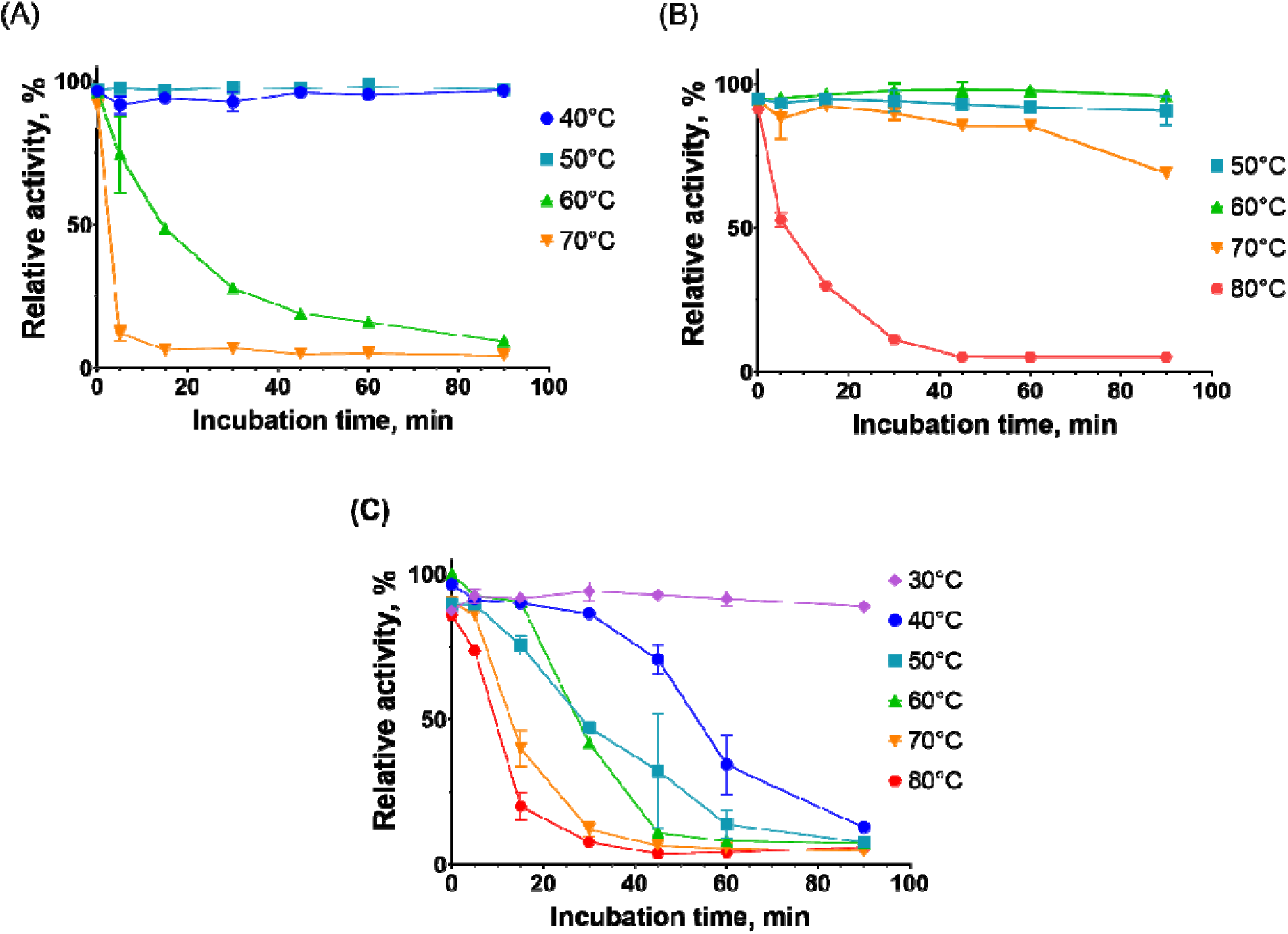
Thermostability of RNAses HII *G. stearothermophilus* (A), *T*. *thermophilus* (B), and *E*. *coli* (C). To assess thermostability, 0.01 U_1_ of each enzyme was incubated for 5–90 minutes at varying temperatures, followed by activity assay. A duplex DNA oligonucleotide with a single rNMP on a fluorescently-labelled strand served as a substrate. All reactions were conducted in the reaction buffer for Eco RNase HII (NEB) at the optimal temperature for each enzyme. Y-axis represents relative activity of RNase HII, and X-axis demonstrates a pre-incubation time. Curve colors correspond to different pre-incubation temperatures as specified in a legend for each panel.

### 1.3. Optimal reaction buffer

Optimal buffer composition allows for more efficient enzyme use, increasing reaction yield and speed. For large-scale practical applications, a reagents cost defines affordability of a technique; thus, using less enzyme makes the method more economical. Additionally, determining the range of working conditions helps to select an optimal buffer for methods assuming simultaneous use of multiple enzymes. For all reaction conditions assessment, we used H2-TM3/H2-TM3RT substrate.

To select the optimal reaction buffer, different cofactors, salts, and pH levels were consecutively tested for each studied RNase HII. First, we evaluated optimal cofactors. Based on previous studies, Mg^2+^, Mn^2+^, and Co^2+^ were chosen as suitable cofactors [5,22]. No significant activity was registered with other divalent cations. The selected cofactors were titrated across a range of 1–15 mM while all other reaction conditions were kept constant. The results of titration are presented in Figure 5. Gst RNase HII demonstrated the highest activity with 5 mM Mg^2+^, Tth RNase HII with 3 mM Mg^2+^, and Eco RNase HII also with 3 mM Mg^2+^. These magnesium concentrations were used in subsequent experiments. The activity of Gst and Tth RNases HII with Mn^2+^ were significantly lower than with Mg^2+^, about 50% and 10%, respectively, and Eco RNase HII was also less active with Mn^2+^. When Co^2+^ concentration exceeded 1 mM, the substrate degraded significantly even without the enzyme present. Therefore, Co^2+^ was excluded from the further analysis, and the respective data are not shown in Figure 5.

**Figure 5.**
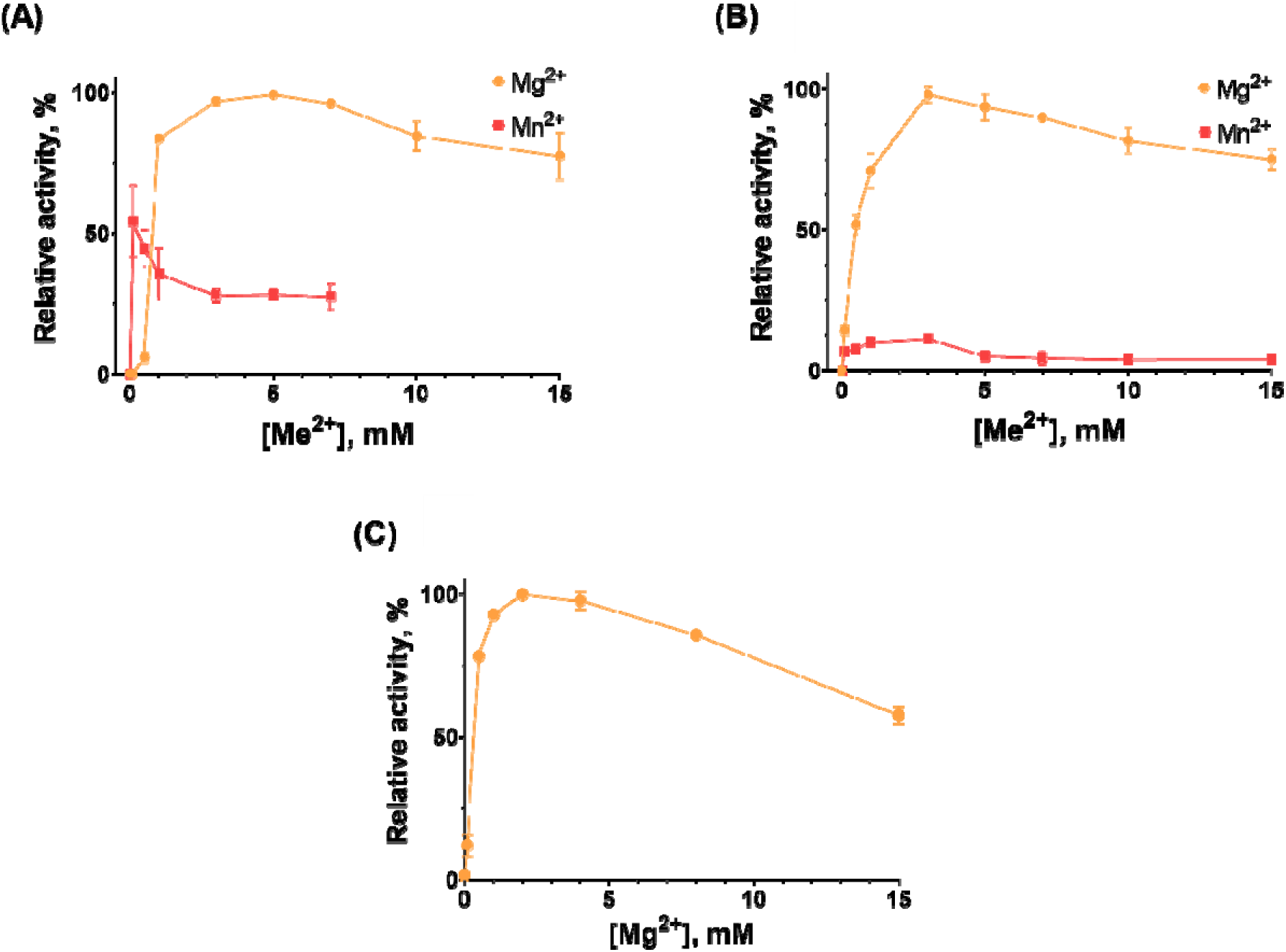
Optimal cofactors for RNases HII *G*. *stearothermophilus* (A), *T*. *thermophilus* (B), and *E. coli* (C). A duplex DNA oligonucleotide with a single rNMP on a fluorescently-labelled strand served as a substrate. Divalent ions were titrated over a range of 0.1–15 mM, while all other reaction conditions were kept constant. The Y-axis represents relative activity of RNase HII, and the X-axis indicates the concentration of divalent cations. Curve colors correspond to cations as specified in a legend for each panel.

Next, we determined the optimal pH by varying this parameter within the range of 5.8–9.5; all other reaction conditions were kept constant. The results of pH optimization are presented in Figure 6. For Gst and Tth RNases HII, the optimal pH was 9.5, and 8.5 for Eco RNase HII. Presumably, the activity of Gst and Tth RNases HII could be higher at a higher pH, but such conditions are unlikely to be optimal for most practical applications. The substrate also degraded noticeably in reaction buffers with pH above 8.0, reaching about 3% degradation at pH 9.5. All subsequent experiments were performed using pH 9, because of relatively high enzymes activity and minimal substrate degradation.

**Figure 6.**
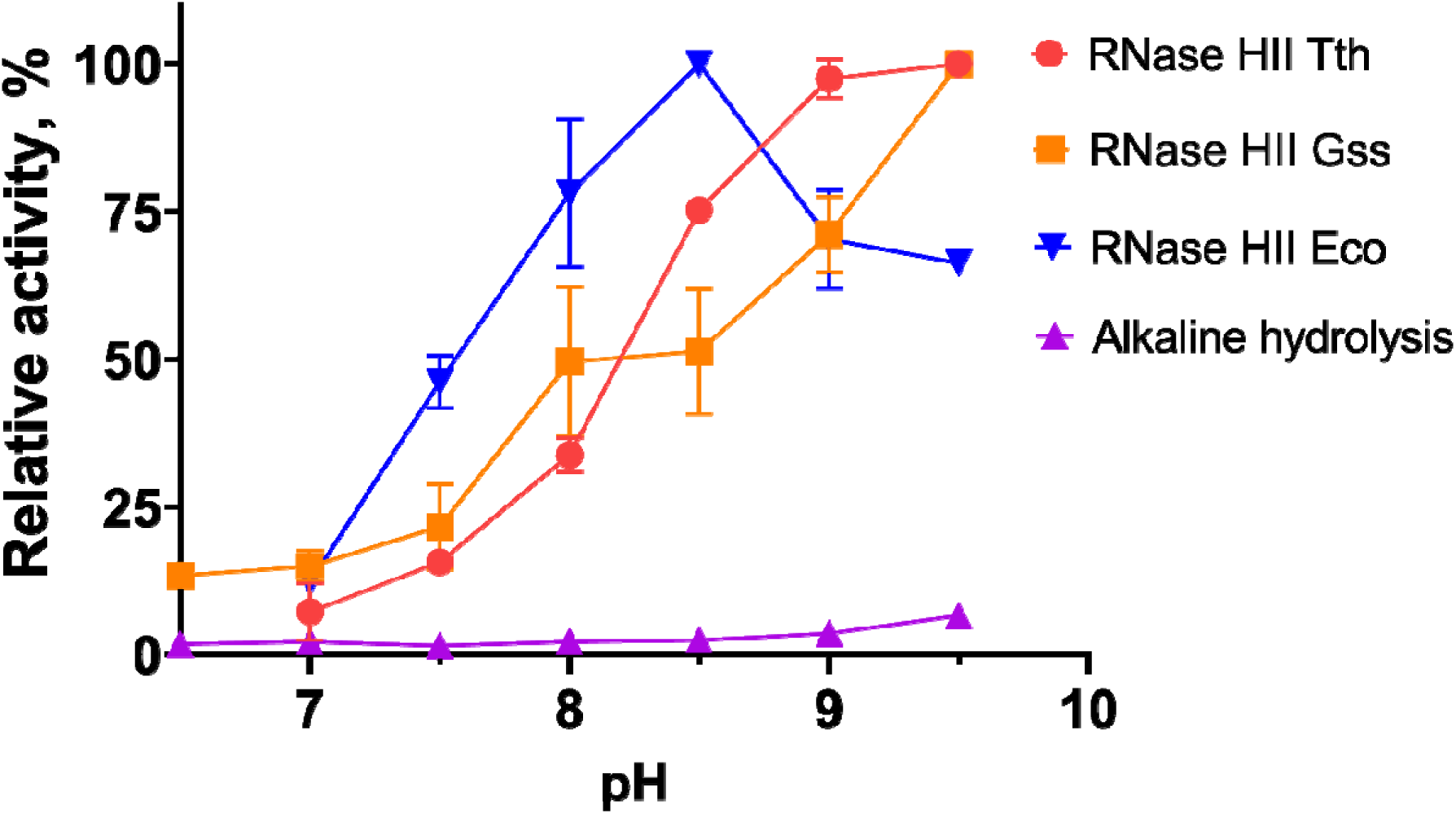
Optimal pH for RNases HII *G. stearothermophilus*, *T. thermophilus* and *E. coli*. A duplex DNA oligonucleotide with a single rNMP on a fluorescently-labelled strand served as a substrate. pH was varied across the range of 5.8–9.5 while all other reaction conditions were kep constant. The Y-axis represents the relative activity of each RNase HII, and the X-axis indicates pH. Curve colors correspond to enzymes as specified in a legend for each panel. The alkaline hydrolysis curve demonstrates substrate degradation without RNase HII.

Finally, the salt composition was optimized. Na^+^, K^+^, and NH_4_ ^+^ were chosen as cations, and Cl^-^, SO4^2-^ served as anions. The activities of RNases HII were tested in the presence of all six salts titrated over a range of 5–150 mM, while other conditions were kept constant. The results of salts titration are presented in Figure 7. Gst RNase HII showed the highest activity with 50 mM NH4Cl, Tth RNase HII with 50 mM KCl, and Eco RNase HII with 25 mM K2SO4. Notably, the impact of salt concentration on enzyme activity was greater than that of the salt type. The activity of all three studied RNases HII decreased with increasing salt concentration in a similar manner, and due to higher ion strength of sulfates, they showed a steeper decrease in activity. Relative activity was also similar across all combinations of cations and anions. The only exception was Gst RNase HII, which was inhibited by Na^+^ and K^+^ cations. Thus, optimal buffer compositions were as follows: 50 mM Tris-HCl pH 8.5, 50 mM NH_4_Cl, 5 mM MgCl_2_ for Gst RNase HII; 50 mM Tris-HCl pH 8.5, 50 mM KCl, 3 mM MgCl_2_ for Tth RNase HII; and 50 mM Tris-HCl pH 8.5, 50 mM NH_4_Cl, 2 mM MgCl_2_ for Eco RNase HII. Although Eco RNase HII have shown slightly greater activity with 25 mM K_2_SO_4_ in our assays, 50mM NH_4_Cl was used chosen as optimal salt type and concentration for this enzyme as it has more consistent activity over a broader concentration range, and such a small difference of 5% lies within the margin of error for our method of measurements. For Gst and Tth RNase HII the difference between highest activity with a given salt was greater than 20% in comparison to the other salt types.

**Figure 7.**
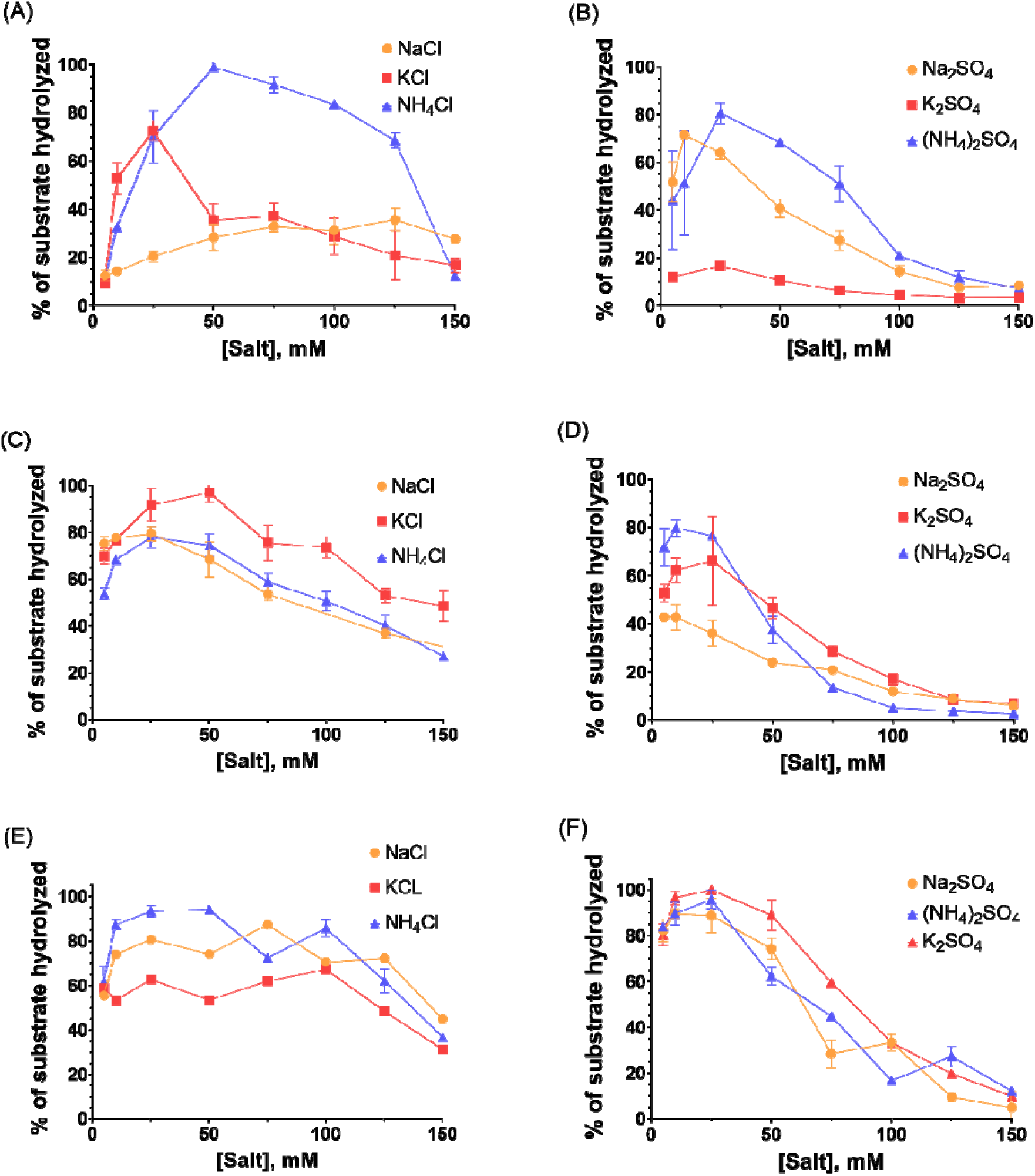
Optimal salts for RNases HII *G. stearothermophilus* (A & B), *T. thermophilus* (C & D), and E. coli (E & F). Panels A, C, and E depict salts with Cl^-^ anion, and panels B, D, and F demonstrate activity with SO^4-^ anions. A duplex DNA oligonucleotide with a single rNMP on a fluorescently-labelled strand served as a substrate. Salts were titrated over a range of 5–150 mM while all other reaction conditions were kept constant. The Y-axis represents the relative activity of RNase HII, and the X-axis indicates the concentration of salt in the reaction buffer. Curve colors correspond to salts as specified in a legend for each panel.

### 1.4. Molar activity and kinetic parameters of RNases HII

After optimization of reaction conditions, the studied RNases were titrated to compare their specific activities under optimal settings. Under optimal conditions, the specific activity of Gst and Tth RNases HII were 1.15- and 2.9-fold higher, respectively, than in the buffer recommended by NEB for Eco RNase HII. The latter buffer was composed for Eco RNase HII, and was almost identical to the buffer we optimized for Eco RNase HII; therefore, the molar activity of this enzyme remained unchanged (Table 2).

**Table 1.**
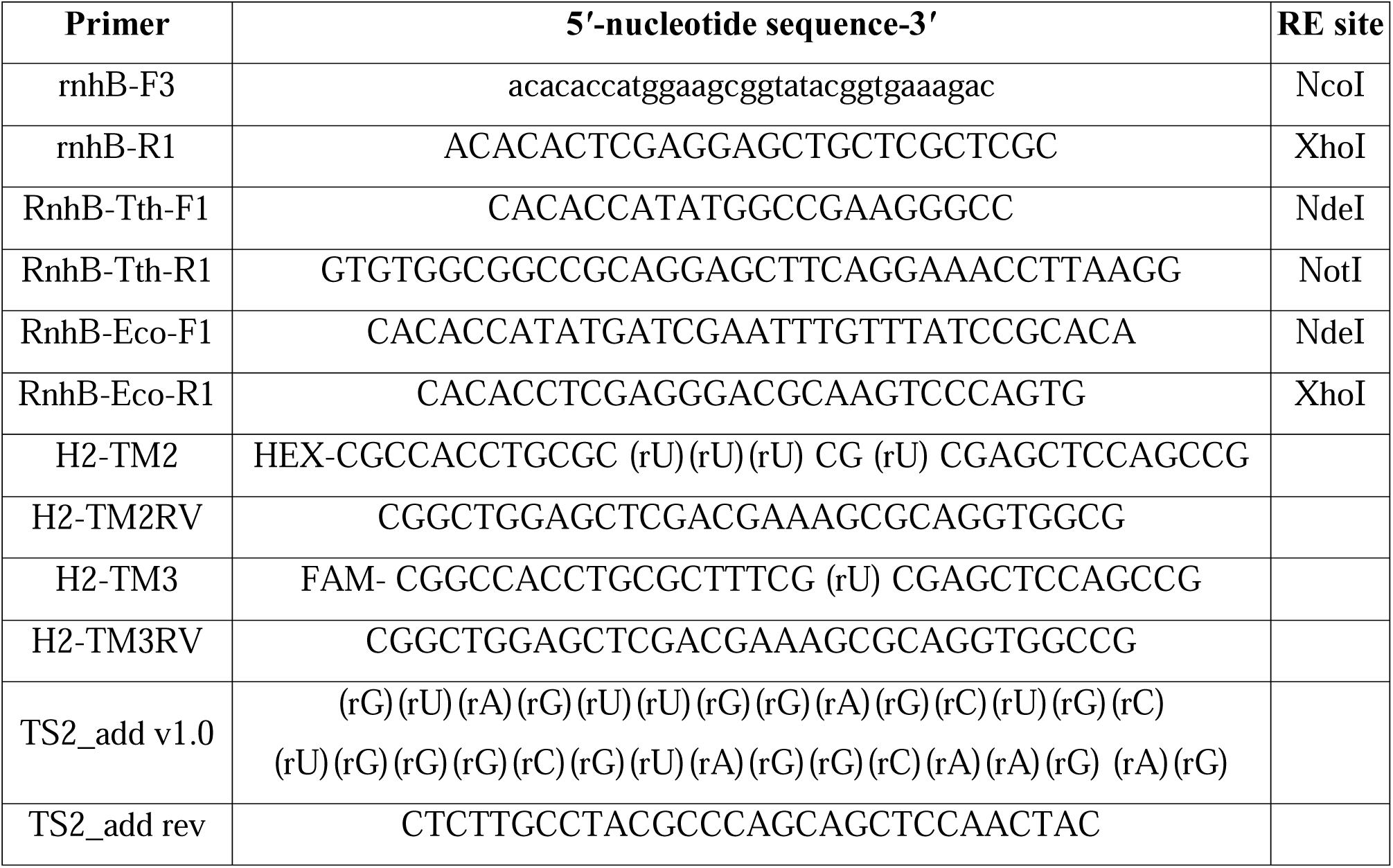
List of oligonucleotide primers and probes.

**Table 2.** Molar activities of RNases HII from different microorganisms.

|  | Gst RNase HII | Tth RNase HII | Eco RNase HII |
| --- | --- | --- | --- |
| Amino acid length | 259 | 203 | 198 |
| Molecular mass, kDa | 28.85 | 21.77 | 21.52 |
| Measured RNase HII molar activity*, U<sub>1</sub> | 31000 ± 3200 | 277000 ± 10500 | 180000 ± 7200 |
| Measured RNase HII molar activity in optimized buffer, U<sub>1</sub> | 36200 ± 1350 | 810000 ± 22600 | 176000 ± 7700 |
| Previously measured RNase HII molar activity*, U<sub>2</sub> | 2.7 ± 0.037 [5] | ~5 [22]; ~10 [14] | - |
| Measured RNase HII molar activity**, U<sub>3</sub> | 74 | 0 | 6670 |
| Previously measured RNase HI molar activity**, U<sub>3</sub> | 20.1 ± 0.9 [5] | 0 [14,22] | ~1500 [22] |
| pI | 8.0 | 7.8 | 7.1 |
| Charge at pH 8.5 | -1.3 | -0.7 | -2.9 |

\* RNase HII activity was measured using the DNA<sub>18</sub>-RNA<sub>1</sub>-DNA<sub>13</sub>/DNA substrate as described above in Material and methods. In the cited papers, different reaction conditions were used, substrates lengths also varied. For Gst RNase HII, the reaction buffer and conditions were significantly different from used in the present work. Notably, the reaction temperature was 30°C, and 50 mM Mg<sup>2+</sup> was used. For Tth RNase HII, different concentrations of salts and BSA were used, and reactions were conducted for 15 minutes at 37°C.
\*\* Our measurements with the RNA/DNA substrate were performed as described above in Material and methods. In cited papers, different reaction conditions were used, substrates lengths also varied. For all RNases HII, we used the same buffer and reaction conditions as for DNA<sub>18</sub>-RNA<sub>1</sub>-DNA<sub>13</sub>/DNA substrate.

Several different approaches have previously been used to determine specific activity of RNases HII. Here, one unit (U_1_) of activity was defined as the enzyme’s amount hydrolyzing 100 pmol of DNA_18_-RNA_1_-DNA_13_/DNA (H2-TM3) substrate after 30 minutes of incubation at optimal temperature, using the buffer recommended by NEB for *E. coli* RNase HII with 0.1 mg/mL BSA. The optimal temperatures were 55, 70, 35°C for Gst, Tth and Eco RNase HII, respectively. Under these conditions, the molar activities were 3.1×10^4^ U_1_/mg for Gst RNase HII, 2.8×10^5^ U_1_/mg for Tth RNase HII, and 1.8×10^5^ U_1_/mg for Eco RNase HII. Permanasari et al [5] defined one unit (U_2_) of Gst RNase HII specific activity as the enzyme’s amount hydrolyzing 1 nmol of the DNA_15_-RNA_1_-DNA_13_/DNA substrate per minute at 30 °C. Under these conditions using the same NEB recommended buffer with BSA and a 30 minutes reaction, specific activity of Gst RNase HII was 103 U_2_/mg which is significantly higher than the 2.7 U_2_/mg reported previously [5]. This discrepancy can be explained by different assay conditions, largely due to reaction temperature and Mg^2+^ concentration. For Tth RNase HII, specific activity was not stated directly; however, after conversion, our measured specific activity (277000 U_1_/mg) was also dramatically higher than previously reported value (∼3000 U_1_/mg) [14]. The same explanations given for Gst RNase HII can be applied here. For Eco RNase HII, the previously used substrate type and the definition of specific activity did not allow comparison of our results with those reported by other groups [14,22,23]. We also defined U_3_ as the enzyme’s amount hydrolyzing 100 pmol of RNA_30_/DNA_30_ under the conditions, identical to the ones described for H2-TM3 substrate. With this substrate we also observed much higher values in comparison with cited papers—74 U_3_ compared to 20.1 U_3_ for Gst RNase HII, and 5.6^10^3^ U_3_ compared to ∼1500 [22] for Eco RNase HII. Tth RNase HII did not cleave this substrate under used reaction conditions (Table 2).

Kinetic parameters of all three studied RNases HII were assessed using the H2-TM3 substrate (Table 3). Among them, Tth RNase HII exhibited the highest catalytic turnover rate, with a k_cat_ of 0.00088 min^-1^, followed by (Eco 0.0312 min^-1^) and Gst RNase HII with the lowest turnover rate at 0.28 min^-1^. The K_m_ values for Gst and Eco RNases HII nearly identical at 0.30 and 0.31 µM, respectively, whereas Tth RNase HII showed the highest K_m_ 0.85 µM.

**Table 3.**
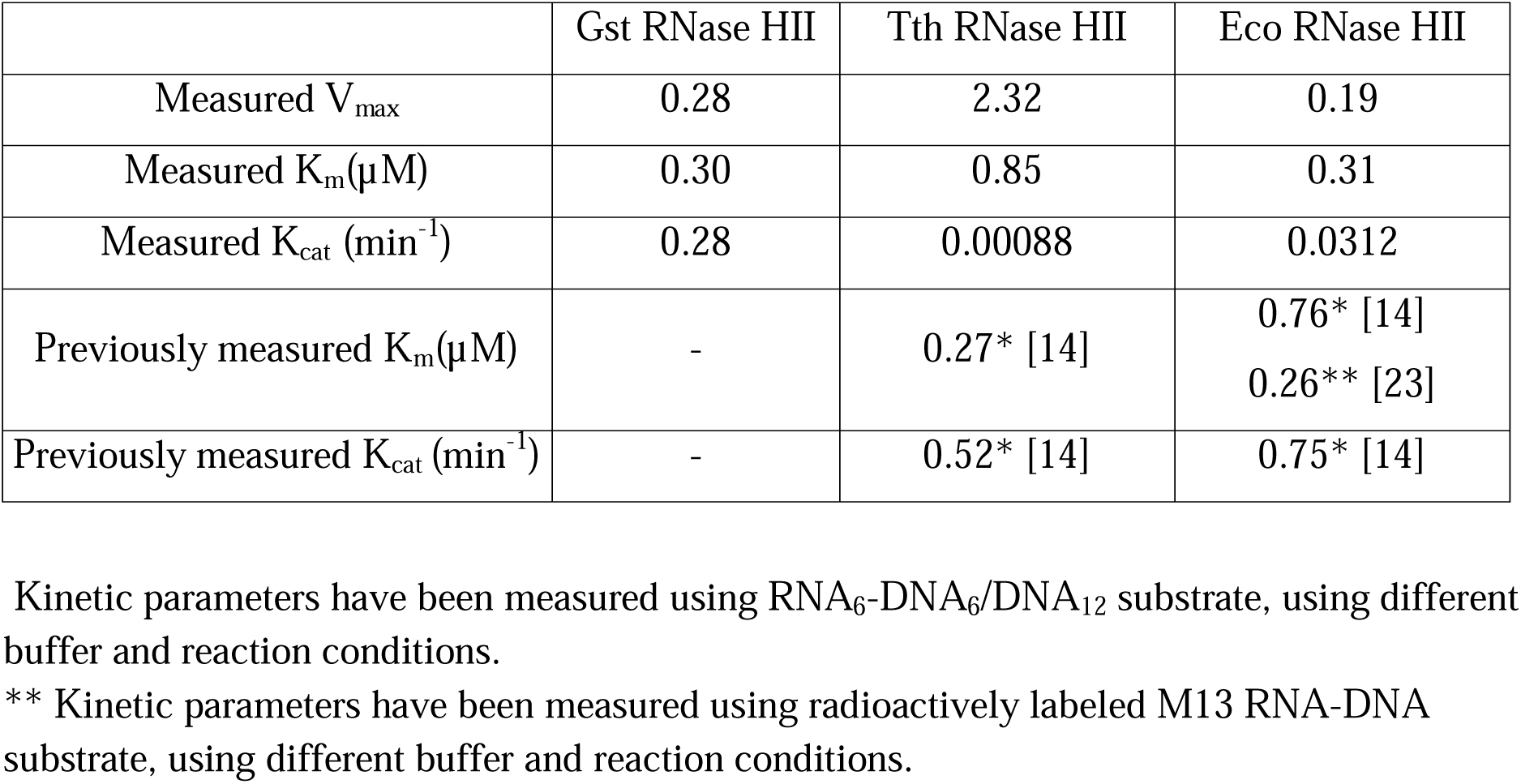
Kinetic parameters of Gst, Tth and Eco RNases HII with H2-TM3 substrate.

Due to differences in substrates and assay conditions employed in the previous studies, direct comparisons with published kinetic data are not feasible. The kinetic values obtained for Tth and Eco RNases HII, along with the substrates used are summarized in Table 3. Kinetic parameters for Gst RNase HII remain limited in the available literature.

### 1.5. Effect of BSA on specific activity of RNases HII

Various substances are often used to enhance efficacy of enzyme’s performance, like betaine, DMSO, trehalose, SSB proteins, and BSA in PCR. These additives can enhance DNA melting [25], mitigate inhibitor’s effect [26], enhance enzyme’s stability [27], etc. One of well-known such substances is BSA [28], a popular stabilizing agent, that prevents thermal inactivation of enzymes, decreases inhibition by unwanted contaminants [29], protects enzymes from non-productive binding or sorption on tube’s walls [30]. In our initial experiments, we observed relatively fast decrease of activity when RNases HII were titrated. Usage of low-binding tubes partially solved this problem, and it made us to supply reactions with BSA to further prevent sorption on tubes. As expected, BSA increased apparent specific activity of all studied RNases (Table 4).

**Table 4.** The effect of 0.1mg/ml BSA on specific activity of RNases HII.

|  | Gst RNase HII | Tth RNase HII | Eco RNase HII |
| --- | --- | --- | --- |
| No BSA, U<sub>1</sub> | 6100 | 63000 | 58000 |
| BSA, U<sub>1</sub> | 31000 | 277000 | 180000 |
| Relative increase | 5.08 | 4.39 | 3.10 |

The same effect was also observed in the thermostability assay. Even a short 5-minute pre-incubation of 0.01U of all tested enzymes (total reaction volume = 10 µL) in optimal reaction buffers at 4 °C dramatically reduced the apparent specific activity, making impossible measurement of thermostability without using BSA as a stabilizing agent. Therefore, all experiments were performed with 0.1 mg/mL BSA. These findings suggest that the tested enzymes adsorb to the walls of reaction tubes. The presence of 0.1 mg/mL BSA resolved this problem, and titration of BSA in the reaction buffer demonstrated a loss of RNase HII activity when the BSA concentration decreased (Figure 8).

**Figure 8.**
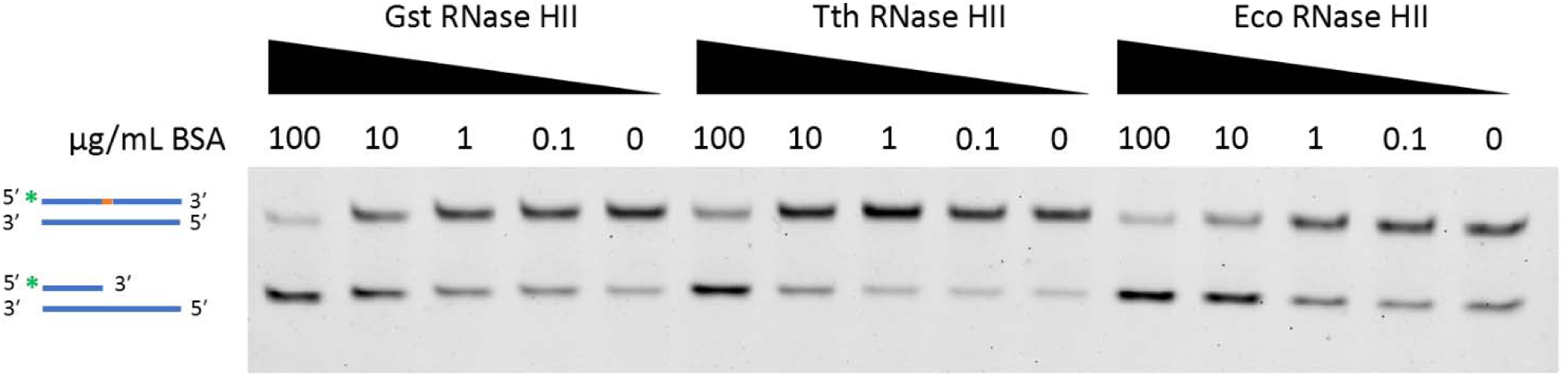
Effect of BSA on RNases HII activity. Each reaction mixture contained 0.01 U_1_ of the specified enzyme. Concentration of BSA is indicated above each lane in µg/mL. The scheme on the left illustrates the substrate (upper band) or the hydrolyzed product (lower band), where blue indicates DNA nucleotides and orange marks an RNA nucleotide. The green asterisk «*» represents the location of FAM on the oligonucleotide.

### 1.6. Substrate specificity

All previously described assays were conducted using the dsDNA substrate with a single rNMP on the FAM-labeled strand (H2-TM3). To test activity of studied RNases HII on other substrates, we used FAM-labeled DNA-RNA_3_-DNA_2_-RNA_1_-DNA (H2-TM2) and oligo(r)NMP_30_ (TS2_add v1.0) oligonucleotides coupled with the respective unlabeled complementary DNA oligonucleotides. The amount of the enzyme with every type of substrate is represented in U_1_, which were calculated for each enzyme in optimal conditions using the buffer recommended by NEB for RNase HII Eco with H2-TM3 substrate. We were able to use more of Gst RNase HII, due to its much higher solubility in the storage buffer. The results of substrate specificity analysis are presented in Figure 9.

**Figure 9.**
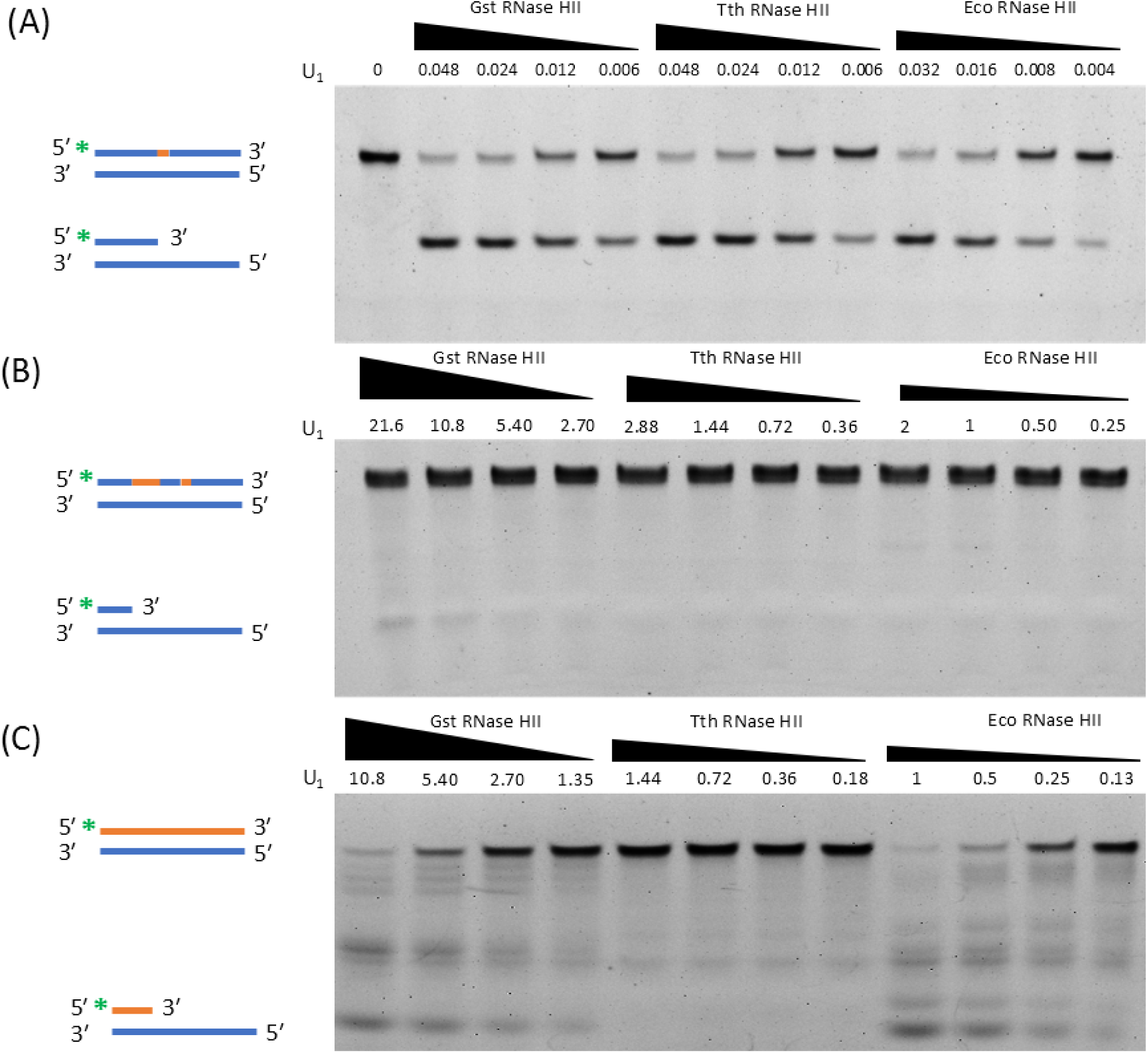
Substrate specificity of Gst, Tth and Eco RNases HII with different substrates. Three substrate types were used: DNA-RNA1-DNA/DNA (A), DNA-RNA_3_-DNA_2_-RNA_1_-DNA/DNA (B), and oligo(r)NMP_30_/DNA (C). Schemes on the left illustrate the substrate and the hydrolysis product, where blue indicates DNA nucleotides and orange marks RNA nucleotides. The green asterisk «*» represents the location of FAM on the oligonucleotide. The amount of each enzyme is indicated in units of activity (U_1_) above each lane, and triangles serve as a visual cue representing relative amounts of RNases HII.

Theoretically, H2-TM2 could be hydrolyzed by either RNases HII or HI, but all three studied RNases HII did not cleave this substrate in the Eco RNase HII (NEB) buffer, with the exception of Gst RNase HII in the highest measured concentration. With 1 mM Mn^2+^ as a cofactor, the results remained the same (data not shown). In contrast, Gst and Eco RNases HII degraded the blunt oligo(r)NMP_30_/DNA duplex. However, their molar activities with this substrate were significantly lower—420-fold for Gst and 27-fold for Eco RNase HII—comparing to the conventional DNA_18_-RNA_1_-DNA_13_/DNA substrate. Additionally, these two enzymes cleft the substrate in different sites. As shown previously [14,22], RNase HII Tth was unable to hydrolyze the RNA/DNA substrate.

## 2. Discussion

Since the discovery of prokaryotic RNases HII, a number of these enzymes have been purified and studied [1,4–14]. However, different methodology does not allow to directly compare results of these researches and use them for deducing reasons behind RNase HII functioning in specific conditions. Even for relatively well-studied enzymes from *E. coli* and *T. thermophilus*, some important aspects are not fully reported including effect of various salts on enzyme’s activity. This lack of empirical data makes difficult the selection of optimal RNase HII for practical applications and reduces their potential usefulness. These enzymes serve as a convenient tool to cleave different types of primers and molecular probes. In probes, the cleavage releases a fluorophore leading to increase of fluorescence. In primers, RNase HII removes a blocked 3-OH end enabling elongation [6]. The latter is useful in DNA amplification methods, as rNMP-containing oligonucleotides must anneal to a complementary DNA for a cleavage by RNase H. Thus, specificity of simplification is increased, because non-specific elongation of primers is blocked. Fluorescently-labelled probes allow to monitor results of isothermal amplification (LAMP) in a real-time manner [31]. In this paper, we chose three already studied and used RNases HII from different organisms and compared their performance in various conditions.

Surprisingly, expression of heterologous RNases HII in *E. coli* cells was inefficient, while recombinant *E. coli* RNases HII was produced in a high amount. In the last case, the yield was up to 50 mg of recoverable protein per gram of biomass, whereas for Tth RNase HII, less than 0.1 mg of enzyme per 1 gram of biomass was achieved. The recombinant Tth RNase HII was almost invisible in SDS-PAGE, but RNases HII activity was detected in post-induction probes with no RNase HII activity detected in pre-induction samples. Different codon usage bias is an often reason behind inefficient production of heterologous proteins. However, expression of Gst and Tth RNases HII in the Rosetta 2 (DE3) strain with the pRARE2 plasmid, coding tRNAs rare in *E. coli*, was also disappointing and similar to the previously used BL21 (DE3) pLysS strain. This observation hints on the presence of a compensatory mechanism mitigating possible negative effect of RNase HII hyperexpression that is specific only for the innate *E. coli* enzyme but not for its counterparts from other organisms.

Another problem of RNases HII purification was a low solubility of recombinant enzymes. We managed to recover less than 0.1 mg of Tth RNase HII and could not evaluate its solubility, but purified in a rather large amount, Eco RNase HII dissolved only in concentrations lower than ∼0.5 mg/ml. This solubility was achieved only by addition of 2 M urea to the lysis, chromatography and storage buffers. It should be noted that urea was diluted to less than 1 mM in final reaction buffer and minimally affected the observed enzyme’s activity. A high concentration of NaCl—600 mM—in the storage buffer increased solubility of Eco RNase HII, but it could affect analysis of RNase HII activity at low salt concentrations. Thus, the storage buffer was supplied with 150 mM NaCl. pH of the storage buffer (7.5-9.5) did not significantly affect Eco RNase HII solubility. Regarding Gst RNase HII, the enzyme dissolved up to 4 mg/mL immediately after chromatography. However, after prolonged storage in 55% glycerol buffer, a substantial amount of enzyme precipitated, and the concertation of a the stable Gst RNase HII solution was only 1.1 mg/ml. Taking into mind a positive effect of BSA on RNase HII activity, we assumed the stabilization of RNase HII in vivo by proteins, substrates or other agents which is not possible in vitro.

During assessment of optimal temperature, the observed optimal temperature was lower—5–10 °C—than expected for Gst and Tth RNases HII based on growth temperature of the respective hosts. Plausibly, protein-protein interactions are also responsible for this discrepancy. Previously, *Archaeoglobus fulgidus* RNase HII, possessing a PIP-box, was reported to form a complex with the cognate PCNA. The addition of Afu PCNA increased the Afu RNase HII processivity on different substrates, including DNA-RNA_1_-DNA/DNA [32]. Similar protein-protein interaction could explain lower temperature optimums and lower than expected thermostability of Gst and Tth RNases HII.

As described in section 3.2.3, our measurements of molar activity were significantly different to that of previously reported. This discrepancy can be attributed to differences in methodology, e.g., buffer composition, reaction temperature, or substrate length and sequences were different in each study. Notably, in these studies, the BSA concentration in reaction buffers varied from 50 to 10 µg/ml, and, as we showed in 3.2.3, BSA can have an effect on apparent specific activity.

Eco and Gst RNases HII were able to hydrolyze RNA/DNA substrate, e.g., possess RNase HI activity, while Tth RNase HII cleft only the DNA-RNA_1_-DNA/DNA substrate. The same substrate preferences were previously reported by Ohtani et al. studied activities of Eco, Bsu and Tth RNases HII on different RNA-DNA/DNA duplexes [22]. It should be also noted that Gst and Eco RNases HII also demonstrated different hydrolysis patterns as well as Bsu and Eco RNases HII. Yet, the DNA-RNA_3_-DNA_2_-RNA_1_-DNA/DNA substrate remained intact, all studied RNases were unable to cleave this type of DNA-RNA-DNA/DNA duplex. This observation rises question about the number of DNA nucleotides on the 5′-end of a substrate that is sufficient for recognition of the duplex by RNases HII. The occurrence of complex DNA-RNA-DNA-RNA-DNA substrates in vivo seems to be low because of a relatively low rate of RNA nucleotides incorporation in during replication of *E. coli* genome (close to 1 rNMP per 2,300 dNMPs). Plausibly, repair of these misincorporation events involves RNase HI, cleaving RNA-DNA/DNA duplexes with multiple rNMPs.

Several limitations of the presented study should be listed. First, we compared biochemical properties of only three RNases HII, but comprehensive analysis of structure-function relation and uncovering reasons of different substrate specificity require a broader set of enzymes. Second, Tth RNAse HII could be affected by contaminant proteins remained after purification of this enzyme. Third, the substrate melting temperature was close to 75°C. The optimal temperature of Tth RNase HII could exceed 80°C, and the H2-TM3 substrate is unfitting for such temperatures. It should be noted that in all studies concerning Tth RNase HII [14,22], the optimal temperature was not determined. Thus, the true optimal temperature of Tth RNase HII can actually be higher than our observation. Fourth, poor solubility and low expression level did not allow us to assess substrate binding by EMSA and thermostability by differential scanning fluorimetry (DSF).

## 3. Conclusions

In the present study, we compared biochemical properties of previously cloned RNases HII from *E. coli, Geobacillus stearothermophilus and Thermus thermophilus*. Optimal reaction conditions—temperature, pH, salts, divalent cofactors—were determined; thermal stability and specific activity were also assessed. As expected, the most thermostable enzyme was Tth RNase HII, and Mg^2+^ was the most preferential cofactor for all RNases. Taking into account alkaline hydrolysis of RNA, the studied RNases HII were the most active at pH 9.0. Gst RNase HII was partially inhibited by K^+^ ions, while other enzymes did not demonstrate preferences for any salt. JRNase activity of Tth RNase HII was confirmed, while the enzymes from *E. coli* and *G. stearothermophilus* exerted typical activities for RNases HII. All three RNases did not cleave a DNA-RNA_3_-DNA_2_-RNA_1_-DNA/DNA substrate. The presented results will facilitate usage of RNases HII in practical applications and provide a basis for further studies of RNases HII from diverse organisms.

## 4. Materials and Methods

### 4.1. Cloning of *G. stearothermophilus*, *T. thermophilus* and *E. coli* RNases HII

The coding sequences of *rnhB* genes from *E. coli*, *G. stearothermophilus*, and *T. thermophilus* were amplified using RnhB-Eco-F3/R1 primers, RnhB-Gst-F3/R1, rnhB-Tth-F1/R2, (Table 1) with NdeI/XhoI, NcoI/XhoI, NdeI/NotI restriction sites, respectively, allowing the in-frame ligation into the pET23d (Gst) or pET23a (Tth and Eco) vectors (Novagen, Darmstadt, Germany). PCR was carried out using *E. coli*, *G. stearothermophilus*, and *T. thermophilus* genomic DNA as templates. The resultant DNA fragments and the vectors were digested with the respective restriction endonucleases (SibEnzyme, Novosibirsk, Russia) and ligated using 100 units of T4 DNA ligase (Biosan, Novosibirsk, Russia). The ligation mixes were used to transform competent cells of the *E. coli* XL1-Blue strain (Stratagene, CA, La Jolla, USA). Absence of mutations in the resulting plasmids was proved by Sanger sequencing performed on ABI 3130XL GeneticAnalyzer (Applied Biosystems, MA, Bedford, USA) using BigDye 3.1 kit (Genomics Core Facility, ICBFM SB RAS, Novosibirsk, Russia). The resulting plasmids named pRnhB-Eco, pRnhB-Gst, pRnhB-Tth were purified from night cultures in 50 mL of LB using QIAGEN Plasmid Midi Kit (Qiagen, Venlo, Netherlands) in accordance with the manufacturer’s protocol.

### 4.2. Expression and purification of *G. stearothermophilus*, *T. thermophilus* and *E. coli* RNases HII

A starter culture of *E. coli* Rosetta2 (DE3) strain (BL21 (DE3) pLysS for Tth and Eco) (Novagen, Darmstadt, Germany) harboring the plasmid pRnhB-Gst (pRnhB-Tth and pRnhB-Eco) was grown to OD600 = 0.3 in LB with 100 μg/mL ampicillin at 37 °C. In a LiFlus GX fermenter (Biotron Inc., Bucheon, South Korea), 4 L of LB with 100 μg/mL ampicillin were inoculated with 20 mL of the starter culture, and the cells were grown to OD600 = 0.6 at 37 °C. The expression of RNases was induced by adding IPTG up to 1 mM concentration. After induction for 4 h at 37 °C, the cells were harvested by centrifugation at 4000× g and stored at −70 °C.

For protein purification, the cell pellet (2.5 g) was resuspended in 20 mL of a lysis buffer (50 mM Tris-HCl pH 8.0, 300 mM NaCl, 1 mM PMSF, 1 mg/mL lysozyme) and incubated for 30 min on ice followed by sonication. After lysis, the soluble fraction was separated by two consequent centrifugation steps at 20,000× g for 30 min. Soluble proteins were precipitated ON by 60% saturation with (NH4)2SO4, followed by centrifugation at 10,000× g for 30 min. The resulting pellet was suspended in lysis buffer without lysozyme and loaded onto a 2 mL IMAC column (Bio-Rad, Hercules, CA, USA) pre-equilibrated with buffer A (50 mM Tris-HCl pH 8.0, 0.3 М NaCl), followed by washing the column with 25 mL of buffer A with 1 M NaCl. Bound proteins were eluted using 10 column volumes and a 0–100% linear gradient of buffer B (buffer A with 0.5 M imidazole). After affinity chromatography, the fractions with Gst RNase HII were pooled, diluted 1:10 with buffer C (20 mM Tris-HCl, 10 mM NaCl, pH 8.0) and loaded onto a column with 10 mL of Heparin SepFast Media (BioToolomics, Consett, UK) pre-equilibrated with buffer C. The column was washed with 50 mL of buffer C, and bound proteins were eluted by 30 column volumes and a 0–100% linear gradient of buffer D (20 mM Tris-HCl, 1 M NaCl, pH 8.0). The fractions with Gst RNase HII were pooled, dialyzed against a storage buffer (25 mM Tris-HCl, 150 mM NaCl, 0.1 mM EDTA, 50% glycerol, pH 8.0, 1 mM DTT), and stored at −20 °C. All the fractions from each step were analyzed using SDS-PAGE. The electrophoretic purity of the isolated Gst RNase HII was not less than 95%. The concentration of purified Gst RNase HII was measured using a standard Bradford assay.

Tth and Eco RNases HII were expressed and purified in a similar way up to affinity chromatography. After affinity chromatography, fractions of Tth and Eco RNases HII were loaded on MacroPrep DEAE Support (Bio-Rad, Hercules, CA, USA) using the same buffers and protocol as for Heparin SepFast Media and Gst RNase HII. During ion-exchange chromatography, the majority of the enzymes did not bind to the DEAE resin and remained in flowthrough. RNases were precipitated by 45% saturated solution of (NH4)2SO4, followed by centrifugation at 20,000× g for 20 min. Precipitates were dissolved in the minimum amount of the storage buffer (25 mM Tris-HCl, 150 mM NaCl, 0.1 mM EDTA, 50% glycerol, pH 8.0, 1 mM DTT) and stored at −20°C. The electrophoretic purity of RNase Tth HII was not less than 80%, and 95% for Eco RNase HII.

All enzymes have been tested for residual nuclease activity by incubation with linear DNA (genomic DNA of T7 phage), supercoiled DNA (pQE30 plasmid) and total RNA from human white blood cells from healthy donors from for 24 hours at 4°C and 37°C. No nuclease activity has been detected.

### 4.3. RNase HII activity assay

The specific activity of RNases HII was assessed by using a 5’-HEX-labeled oligonucleotide React 3 with a single ribonucleotide annealed to a complementary oligonucleotide React 7 at a 1:4 ratio (Table 1). The 10 µL reaction mixture contained 2 pmol of substrate (concentration defined by the labeled oligonucleotide), 50 mM Tris-HCl (pH 8.5), 3 mM MgCl_2_, 75 mM KCl, 5 mM DTT. Reactions were initiated by adding the substrate and immediately transferred to a preheated thermal block for incubation: at 50°C for Gst and Tth RNase HII, and 37°C for Eco RNase HII, for 30 minutes. After incubation, reactions were quenched by adding 10 µL of formamide containing bromophenol blue dye and heating at 95°C for 5 minutes. Reaction products were analyzed by denaturing polyacrylamide gel electrophoresis (PAGE) using a 20% acrylamide gel with bis-acrylamide (30:1), 7 M urea and 0.5× TBE buffer with pre-heating to 55°C. Gels were documented using GelDoc Go (Bio-Rad, USA) and analyzed using Image Lab 6.0 software (Bio-Rad, USA).

Similarly, HEX-labeled oligonucleotide H2-TM2, containing an r(X)-r(X)-r(X)-XX-r(X) motif, and FAM-labeled RNA/DNA substrates were used to test the activity of studied the RNases HII. No hydrolysis of H2-TM2 was detected under any tested conditions, and RNA/DNA substrate was degraded only by Eco and Gst RNases HII, but not by Tth.

### 4.4. Quantification of RNase HII activity and optimal conditions assays

On the electropherograms of activity assays, each lane contained one or two bands: the upper band represented the intact substrate and the lower band corresponded to the digested product. Quantitative analysis was performed using Image Lab 6.0 software in the integrated “Volume Tools” mode. Individual bands were enclosed within rectangles; an additional rectangle was placed between the upper and lower bands to define background, and the background subtraction mode was set to “global”. For intensity measurement, the “Adj. Vol. (int)” was selected, and the intensity value for each band was used in subsequent analysis. For each lane, the product-to-substrate ratio was calculated using an equation А = (1 - (X)/(X+Y)) × 100%, where A is the product-to-substrate ratio, X represents the intensity of the upper band (substrate), and Y represents the intensity of the lower band (product). The resulting value A represents the percentage of hydrolyzed product. The A values for all lanes were visualized using GraphPad Prism 8.0 software, and molar activities were determined for all three RNases HII. One unit (U_1_) of activity was defined as the amount of enzyme required to hydrolyze 100 pmol of H2-TM2/ H2-TM2RV after 30 minutes of incubation at its optimal temperature in a buffer recommended by NEB for Eco RNase HII with 0.1 mg/mL BSA. Similarly, U_3_ was defined, using TS2_add v1.0/ TS2_add rev substrate.

### 4.5. Thermal stability, optimal temperature, pH, ion concentration, divalent cofactors and kinetic parameters

In the following experiments, all conditions were identical to the activity assay, except for the parameter under study. In all reactions, the amount of an RNase HII was 0.01 U_1_. The thermal stability of RNases HII was determined by heating the enzymes, followed by the activity assay. Before activity measurement, reactions without the substrate (8 µL each) containing 0.1mg/ml BSA and 0.01 U_1_ of RNases HII were incubated at 30–80 °C for 5–90 min. The temperature optimum of RNases HII was determined by measuring specific activity at the temperature range of 25–75°C. Optimal pH was evaluated by conducting activity assay at pH ranging from 6.5 to 9.5. Optimal ion and cofactor concentrations were assessed by varying salts and their concentrations: 5–150 mM of NH_4_Cl, KCl, NaCl, Na_2_SO_4_, K_2_SO_4_, (NH4)_2_SO_4_, or 1–15 mM MgCl_2_, 1–15 mM MnCl_2_, 1–15 mM CoCl_2_.

Kinetic parameters were determined by conducting reactions for 2 minutes using varying concentrations of substrate H2-TM3 ranging from 2µM to 0.02µM. Reactions were performed in the buffer recommended by NEB for Eco RNase HII, supplemented with 0.1 mg/mL BSA, and employed 0.1 U_1_ of each enzyme at its optimal temperature in optimized buffer. The amount of hydrolyzed substrate was calculated following the same procedure as in previous assays. K_m_ and V_max_ were computed using standard Michaelis-Menten non-linear regression in Graph prism 8.0.1 software.

## Acknowledgements

This work was supported by the Russian state-funded project for ICBFM SB RAS (grant number 125012300657-2).

## Author Contributions

MLF conceived the study. AAV and IPO designed experiments; AAV, LMN, and MEV performed experiments. AAV and IPO analyzed experimental data. AAV drafted the manuscript and prepared figures; IPO wrote portions of the manuscript. OIP supervised the study; MLF revised the manuscript. All authors reviewed the results and approved the final version of the manuscript.

## Data availability

The data that support the findings of this study are available from the corresponding author upon reasonable request.

## Abbreviations

cDNA: complementary DNA
dNMP: deoxyribonucleotide monophosphate
JRNase: junction ribonuclease
RNA-seq: RNA sequencing
rNMP: ribonucleotide monophosphate

